# Intrinsic brain connectivity after partial sleep deprivation in young and older adults: results from the Stockholm Sleepy Brain study

**DOI:** 10.1101/073494

**Authors:** Gustav Nilsonne, Sandra Tamm, Johanna Schwarz, Rita Almeida, Håkan Fischer, Göran Kecklund, Mats Lekander, Peter Fransson, Torbjörn Åkerstedt

## Abstract

Sleep deprivation has been reported to affect intrinsic brain connectivity, notably reducing connectivity in the default mode network. Studies to date have however shown inconsistent effects, in many cases lacked monitoring of wakefulness, and largely included young participants. We investigated effects of sleep deprivation on intrinsic brain connectivity in young and older participants. Participants aged 20–30 (*n*=30) and 65–75 (*n*=23) years underwent partial sleep deprivation (3h sleep) in a cross-over design, with two 8-minutes eyes-open resting state functional magnetic resonance imaging (fMRI) runs in each session, monitored by eye-tracking. We assessed intrinsic brain connectivity using independent components analysis (ICA) as well as seed-region analyses of functional connectivity, and also analysed global signal variability, regional homogeneity, and the amplitude of low-frequency fluctuations. Sleep deprivation caused increased global signal variability. In our study, changes in investigated resting state networks and in regional homogeneity were not statistically significant. Younger participants had higher connectivity in most examined networks, as well as higher regional homogeneity in areas including anterior and posterior cingulate cortex. In conclusion, we found that sleep deprivation caused increased global signal variability. We speculate that this may be caused by wake-state instability.

## Introduction

### Background

Sleep problems are a public health issue affecting about one third of the general population, of which about one in three reports serious sleep problems^1,2^. Impaired or shortened sleep is a risk factor for mortality and for a number of diseases^3^, as well as accidents^4^. These risks appear to be mediated by impaired biological restoration/recovery^3^. Effects of sleep loss include an increasing number of short lapses of attention (microsleeps)^5^, as well as increased levels of EEG alpha and theta activity; slow, rolling eye movements; and increased subjective sleepiness^6^.

Sleepiness is not well defined in terms of functional brain activation. Early studies using total sleep deprivation have shown reduced glucose uptake in large areas of the brain, including prefrontal and parietal areas^7^. Intrinsic connectivity, referring to functional connectivity in the resting state, i.e. when participants are not presented with any changing stimuli, has been investigated in several earlier studies. We have identified 21 earlier reports, based on 14 unique datasets^8–28^. An in-depth review of this earlier literature is beyond the scope of this paper, and for this reason we have recently made available online an overview of these reports^29^. A consistent and robust finding is that sleep deprivation caused reduced connectivity within the default mode network and reduced anticorrelation to the task-positive network^8,10,22^. Other findings include increased regional homogeneity (ReHo) in different brain areas following sleep deprivation^9,18^, changes in connectivity between the thalamus^12^ and amygdala^13,14^ and cortical areas. We formulated a number of hypotheses to try to replicate these findings (see below).

It is an interesting question whether reduced connectivity, observed in previous studies, is associated with subjective sleepiness, which is a sensitive indicator of sleep loss^30^. There is also a question whether participants in previous studies were able to consistently maintain wakefulness during resting state scanning, not least in light of findings by Tagliazucchi and Laufs^31^, showing that about 50% of non-sleep-deprived participants fall asleep within 10 minutes of resting state imaging. When the fMRI-based classifier developed by Tagliazucchi was tested on data from a sleep deprivation experiment with scanning with eyes open, the probability of being awake was estimated at below 50% for the duration of scanning^23^. Prior studies of intrinsic connectivity after sleep deprivation have mainly relied on self-report to rule out episodes of falling asleep during scanning^29^. The role of age has been investigated in only one study of sleep deprivation and connectivity, which compared sleep deprivation in young participants to an openly available dataset with older participants who had not been sleep deprived^27^. It appears that sleepiness is reduced in older individuals both when sleeping normally and after sleep deprivation^32^. Contrary to intuition, younger individuals seem more susceptible to sleep loss in terms of physiological and self-reported sleepiness^32^. Furthermore, sleep duration decreases with age^33^ and the ability to produce sleep under optimal conditions also falls with age^34^, suggesting that sleep need is reduced. Functional connectivity is generally lower in older compared to younger humans, possibly due to structural changes including white matter degeneration of vascular genesis^35–37^.

The fMRI global signal during rest has been the topic of controversy as a nuisance regressor. Murphy et al.^38^ reported that global signal regression may introduce artifactual anti-correlations between resting state networks (see also Weissenbacher et al.^39^). Saad et al.^40^ reported that global signal regression may also introduce bias in comparisons between groups and time-points, and Guo et al.^41^ reported that global signal regression reduced re-test reliability of salience network estimation in 29 older adults scanned with an interval of a year. In this context, it deserves to be mentioned that the results from the studies that first showed negative correlations between resting state networks, primarily between the default mode and task-positive networks^42,43^ have subsequently been reproduced without use of global brain mean signal regression (e.g. ^44,45^). However, recently, the global signal has become a focus for interest in itself. Schölvinck et al.^46^ reported a correlation between spatially widespread BOLD signal changes and spontaneous fluctuations in local field potentials in monkeys, mainly in the high gamma frequency band. Furthermore, Wong et al.^47^ reported that global signal variability was decreased by caffeine and furthermore that global signal variability was negatively correlated to EEG measures of vigilance^48,49^. In addition, Fukunaga et al.^50^ found an increased BOLD signal amplitude in those individuals that were asleep at the end of a resting state session. These findings suggest that global signal variability is higher in sleepiness. The effect of sleep deprivation on global signal variability has been investigated in one previous report, which reported increased variability after sleep deprivation, but which did not provide inferential statistics for this finding^22^.

The role of prior sleep, apart from sleep deprivation, may also be of interest in relation to resting state connectivity. Killgore et al.^51^ showed that self-reported shorter sleep duration prior to imaging was associated to reduced connectivity. No studies to date have investigated the contents of prior sleep with polysomnography, the gold standard for sleep measurement. Both sleep fragmentation^52^ and suppression of N3 (“deep”) sleep^32^ increase physiological sleepiness. Partial sleep deprivation (PSD) allowing 2 hours of sleep strongly increase physiological and self-reported sleepiness, while PSD allowing 4 hours of sleep only gives marginal increases^53^. Thus, it is an interesting question whether sleep reduction to < 4h causes reduced resting state connectivity.

This study investigated intrinsic brain connectivity measures and global signal variability after partial sleep deprivation and in relation to PSG and reported sleepiness in the Stockholm Sleepy Brain Study 1, a brain imaging study including younger (20-30 years) and older (65-75 years) healthy volunteers.

### Aims

We aimed to investigate the effects of partial sleep deprivation on intrinsic brain connectivity. Specifically, we hypothesised that sleep deprivation would cause

- decreased connectivity within the default mode (DMN), salience, frontoparietal attention, and executive control networks.
- decreased anticorrelation between DMN and the task-positive network (TPN) during resting state.
- changes in thalamocortical connectivity.
- changes in connectivity from amygdala, specifically decreased connectivity between amygdala and prefrontal cortex.
- changes in regional homogeneity
- increased global signal variability

Furthermore, we hypothesised that the above-mentioned measures would correlate to PSG measures of sleep and to self-rated sleepiness, and that older participants would have lower functional connectivity and be less sensitive to sleep deprivation. We also exploratively investigated the amplitude of low-frequency fluctuations (ALFF).

## Materials and methods

### Study design

The study was a cross-over comparison between 3 h partial sleep deprivation and full sleep. This particular sleep duration was chosen because it appears that clear effects on physiological and subjective sleepiness require < 4 h of total sleep time (TST)^53^, because it is highly relevant for sleep problems in the population, and because pilot testing suggested partial sleep deprivation would be less likely to cause participants to fall asleep during the experiment compared to total sleep deprivation. Participants were randomised to undergo both conditions in a counterbalanced order, with an interval of approximately one month. In the interest of ecological validity, participants slept in their own homes in both conditions. Sleep was monitored using ambulatory polysomnography. In the sleep deprivation condition, participants were instructed to go to bed 3 h before the time they would usually get up, and then to get up at that time. MRI imaging was performed in the evening following sleep deprivation or normal sleep (approximately between 18:00 and 21:00), in order to avoid confounding by circadian rhythms^54,55^. Experimenters at the MRI scanner were blinded to participants’ sleep condition.

The project was preregistered at clinicaltrials.gov

(https://clinicaltrials.gov/ct2/show/NCT02000076), with a full list of hypotheses and an analysis plan available on the open science framework (https://osf.io/zuf7t/). Hypotheses were updated in light of new findings reported in the literature after data collection but before data analysis. The study was approved by the Regional Ethics Review board of Stockholm (2012/1870-32). All participants gave written informed consent. Experiments were performed in accordance with the Declaration of Helsinki and applicable local regulations. Methods, data, and technical validation have been reported in detail in a previous manuscript^56^.

### Participants

As described in Nilsonne et al.^56^, participants were recruited by poster advertising on campus sites in Stockholm, on the studentkaninen.se website, and in newspaper ads. Prospective participants were screened for inclusion/exclusion criteria using an online form and eligibility was confirmed in an interview upon arrival to the scanning site. Criteria for inclusion were, first, those required to undergo fMRI procedures and to use the hand-held response box, namely: no ferromagnetic items in body, not claustrophobic, not pregnant, no refractive error exceeding 5 diopters, not color-blind, and right-handed. In addition, participants were required to be 20-30 or 65-75 years old (inclusive), to have no current or past psychiatric or neurological illness, including addiction, to not have hypertension or diabetes, to not use psychoactive or immune-modulatory drugs, to not use nicotine every day, and to have a lower habitual daily caffeine intake corresponding to 4 cups of coffee at most. A further criterion was to not study, have studied, or be occupied in the fields of psychology, behavioural science, or medicine, including nursing and other allied fields. The reason for this was that participants with a background in psychology might try to “see through” the experimental paradigm, whereas participants with a background in medicine may have a less strong emotional response to pictures showing needles or injuries, which were used in two of the experiments not reported in the present paper. The insomnia severity index (ISI)^57,58^, the depression subscale of the Hospital Anxiety and Depression scale (HADS)^59,60^ and the Karolinska Sleep Questionnaire (KSQ)^61^ were used to exclude participants with insomnia symptoms, out-of-circadian sleep patterns, or excessive snoring (see 2.3). For practical reasons, participants were also required to understand and speak Swedish fluently and to live in the greater Stockholm area. Participants were paid 2500 SEK (approx. 280 Euro/360 USD), subject to tax. They were also offered taxi travel to and from the MRI imaging center, in order to avoid traffic incidents following sleep deprivation.

### Sleep measures

As described in Åkerstedt et al. (manuscript in preparation), polysomnography (PSG) recording took place in the homes of the participants using a solid state, portable sleep recorder (Embla). Standard electrode (silver/silver chloride) montage for EEG sleep recording was used (C3, C4 referenced to the contralateral mastoid). In addition, two sub-mental electrodes were used for electromyography (EMG) and one electrode at each of the outer canthi of the eyes were used for electrooculography (EOG). Sleep staging, respiratory and arousal analysis were performed according to the classification criteria of the American Academy of Sleep Medicine (AASM) using the computer-assisted sleep classification system Somnolyzer 24 × 7 (Anderer et al., 2005, 2010). To adapt to AASM scoring, F4 was interpolated. Here the terminology N1, N2, and N3 is used for sleep stages 1-3. Wake within the total sleep period (WTSP) represents time awake between sleep onset and offset and this value is expressed in percent of the total sleep period (TSP). Shifts from any of the sleep stages to wake were expressed as awakenings per hour.

### Experimental task

Two resting state sessions were performed on each of the two visits to the MRI scanner. The first session was in the beginning of scanning, preceded by a 4 min anatomical scan which allowed the participants to acclimatize to the scanner environment. The second session was at the end of scanning, following approx. 1 hour of experiments using emotional stimuli, which will be reported elsewhere. Participants were instructed to look at a fixation cross presented against a gray background, presented using goggles (NordicNeuroLab). During scanning, participants were monitored by eye-tracking. In case of eye-closures of more than approx. 5 seconds, the MRI operator spoke a wake-up call through the participant’s headphones. This happened only once in one participant during resting state scanning among all of the included participants. In the first run, the resting state acquisition period lasted for 8 minutes with no interruptions. In the second run, also of 8 minutes, participants were asked to rate their sleepiness every 2 minutes with the Karolinska Sleepiness Scale (KSS)^6,30^. This is a single-item question with 9 ordinal anchored response alternatives.

### MRI acquisition

We used a General Electric Discovery 3 T MRI scanner. Echo-planar images were acquired using the following settings: flip angle 75, TE 30, TR 2.5 s, field of view 28.8 cm, slice thickness 3 mm, 49 slices, interleaved bottom → up. T1-weighted anatomical scans were acquired with a sagittal BRAVO sequence, field of view 24 cm, slice thickness 1 mm, interleaved bottom → up.

### Preprocessing

To remove task-related interference, volumes scanned during KSS ratings in the second run were cut out of the time series, including two volumes before and four after each rating event. This procedure removed spikes in the time-series for visual and motor networks, occurring at the time of ratings (not shown). We note that a consecutive time-series is not an assumption underlying such analyses of connectivity as were performed on these data. Remaining volumes were concatenated and in order to balance data, each series was trimmed down to the lowest number of remaining scans in any one session, which was 163, corresponding to 7 minutes and 20 seconds. Data were preprocessed in SPM12 using the DPARSFA toolbox^62^. Functional images were slice time corrected, realigned, normalized using DARTEL^63^, resampled to 3x3x3 mm voxel size, spatially smoothed with a 6x6x6 mm kernel, detrended, frequency filtered (0.01-0.1 Hz), and regressed on nuisance covariates including six motion regressors and gray and white matter signal using DARTEL-obtained segmentation. Participants with more than 40 volumes (25%) with framewise displacement (FD) ≥ 0.5 mm in any run were excluded from analysis (*n* = 15 out of 68, leaving *n* = 53 for analysis). Remaining volumes with FD ≥ 0.5 mm were cut after nuisance regression and interpolated using cubic splines. Unless otherwise specified, these data were used for the subsequent analyses. To further reduce the risk of motion confounding, FD parameters were carried forward as regressors of no interest in 2^nd^ level analyses.

### Independent component analysis (ICA)

ICA was performed using the GIFT toolbox^64^. Independent components were estimated using the Infomax algorithm and the ICASSO approach. Following back-reconstruction using spatio-temporal regression, components of interest were extracted for each subject. 2^nd^ level analyses were performed in SPM12 within a general linear model (GLM) framework.

We performed two sets of ICA analyses. The first set was performed on data preprocessed with frequency filtering (as described above), with a gray matter mask derived from DARTEL, and with 20 components. One reviewer suggested that the analysis should be repeated without frequency filtering, without the gray matter mask, and with 30 components. We report the 30-component ICA as the main model.

In 2^nd^ level analyses, masks for specific networks were used based on a previously published parcellation^65^, illustrated in supplemental Figure 1. We also investigated effects within the whole gray substance mask. We did not use network masks derived from the present data, as this would introduce a risk of overfitting.

### Seed-based and cross-correlation analyses

Seed regions for the default mode network (DMN) and the anticorrelated task-positive network (TPN) were based on a previous report on the effect of sleep deprivation on functional connectivity^10^. Seed regions were 9x9x9 mm cubic centered on coordinates reported by De Havas et al.^10^, see supplemental Table 1. Time courses were compared within each run using the DPARSFA toolbox and the resulting z-statistics were entered into a mixed-effects linear regression model with deviation coding for contrasts in R^66^. In addition, following Shao et al.^12^, we defined a thalamus region of interest (ROI) using the Wake Forest University PickAtlas^67–69^, and following Shao et al.^12,13^ and Lei et al.^14^, we selected right and left thalamus ROI:s as well as separate ROI:s for the superficial, centromedial, and basolateral amygdala, as defined by Amunts et al.^70^, using the Jülich atlas^71–73^. Furthermore, at the suggestion of one reviewer, we investigated default mode network using seed-based methods. To capture the default mode network in a manner that was optimized for the present dataset, we investigated seed regions centered on the peak voxel of the posterior midline hub according to the 30-component ICA (MNI 0, −58, 15). We tested a spherical seed with radius 4.5 mm and a box seed with dimensions 9x9x9 mm. We also tested a set of seed regions defined as those areas where either of the two default mode network components from the 30-component ICA showed *z* > 30.

### Regional homogeneity (ReHo)

Using the DPARSFA toolbox, ReHo was estimated in data preprocessed as described above but without smoothing, with cluster size of 27 voxels (3x3x3) and subsequent smoothing. 2^nd^ level analysis was performed in SPM12 as described above.

### Global signal

Global signal was estimated using the DPARSFA toolbox from data preprocessed as described above including all the steps and with the same gray matter mask generated by DARTEL. For each run, global signal variability was determined as the standard deviation of gray matter signal during the run, following Wong et al.^48,49^. Notably, Wong et al. calculated the same measure but called it amplitude rather than variability, and we used that terminology in the registration of hypotheses. To better approximate a normal distribution, standard deviations were log-transformed. Mixed-effects models were then used in R^66^ to investigate effects of sleep deprivation and age group, as well as correlations to putative covariates, with mean framewise displacement as a covariate of no interest in order to decrease the influence of head motion on estimates.

### Amplitude of low-frequency fluctuations (ALFF)

Amplitude of low-frequency fluctuations (ALFF) and fractional amplitude of low-frequency fluctuations (fALFF) were analysed using the DPARSFA toolbox on preprocessed data after nuisance regression but before scrubbing and interpolation, in order to preserve continuity of the time-series. For the same reason, the second run in each session was not included in these analyses, as these runs had volumes censored. Temporal filtering was not performed before ALFF and fALFF analyses, in order to preserve low-frequency fluctuations for analysis.

### Availability of data and code

Structural and functional imaging data are available at https://openfmri.org/dataset/ds000201/. (Note: provisional links to data during review process are provided in the cover letter). Code for preprocessing and analysis, SPM results objects and tables are available at http://dx.doi.org/10.5281/zenodo.581250. Masks and seed regions are available for visualisation and download at http://neurovault.org/collections/LEWNNZLY/.

## Results

### Participants

The final sample consisted of 53 participants (30 young, 23 old), after excluding 15 due to excessive motion (8 young, 7 old). Participant characteristics are given in Table 1. PSG data for the full sample of participants have been reported in Åkerstedt et al. (in preparation), but are repeated here for those participants finally included in resting state analyses.

**Table 1:** Participant characteristics and sleep measures.

|  | Condition | All | Young | Old |
| --- | --- | --- | --- | --- |
| Sex (F/M) |  | 29/24 | 16/14 | 13/10 |
| Age (median, range) |  | 26 (20-75) | 23 (20-29) | 68 (65-75) |
| Total sleep time, minutes (mean, SD) | Full sleep | 412 (76) | 431 (78) | 389 (68) |
|  | Sleep deprivation | 173 (37) | 184 (37) | 158 (33) |
| REM, % (mean, SD) | Full sleep | 19.5 (6.9) | 19.8 (5.6) | 19.2 (8.3) |
|  | Sleep deprivation | 15.4 (8.8) | 14.4 (6.9) | 16.0 (10.0) |
| N1, % (mean, SD) | Full sleep | 18.8 (10.1) | 14.4 (6.6) | 28.6 (11.1) |
|  | Sleep deprivation | 15.6 (9.0) | 10.8 (5.8) | 20.6 (9.4) |
| N2, % (mean, SD) | Full sleep | 44.1 (7.7) | 42.8 (7.7) | 45.8 (7.5) |
|  | Sleep deprivation | 40.7 (11.8) | 35.5 (9.6) | 45.4 (12.0) |
| N3, % (mean, SD) | Full sleep | 17.5 (9.6) | 23.0 (6.8) | 10.4 (8.0) |
|  | Sleep deprivation | 28.3 (16.0) | 39.3 (9.5) | 18.1 (14.3) |
| ESS (mean, SD) |  | 8.2 (3.8) | 7.4 (2.8) | 9.2 (4.6) |
| ISI (mean, SD) |  | 9.9 (2.0) | 10.1 (2.2) | 9.0 (1.3) |
| KSQ sleep quality index (mean, SD) |  | 5.3 (0.5) | 5.3 (0.5) | 5.2 (0.5) |
| KSQ snoring symptom index (mean, SD) |  | 5.7 (0.4) | 5.6 (0.3) | 5.5 (0.5) |

### Measures of sleep and sleepiness

KSS data for the full sample of participants have been reported in Åkerstedt et al. (in preparation), but are repeated here for those participants finally included in resting state analyses. Effects of sleep deprivation, age group, and time in scanner on KSS ratings was investigated using a mixed-effects regression model. Sleep deprivation caused increased KSS (β = 1.88 [95% CI 1.73, 2.03], *p* < 0.001, Figure 1). Young age group was associated to higher KSS ratings (1.33 [0.67, 2.00], p < 0.001), as was time in scanner (0.76 per hour [0.60, 0.92], p < 0.001, Figure 1).

**Figure 1:**
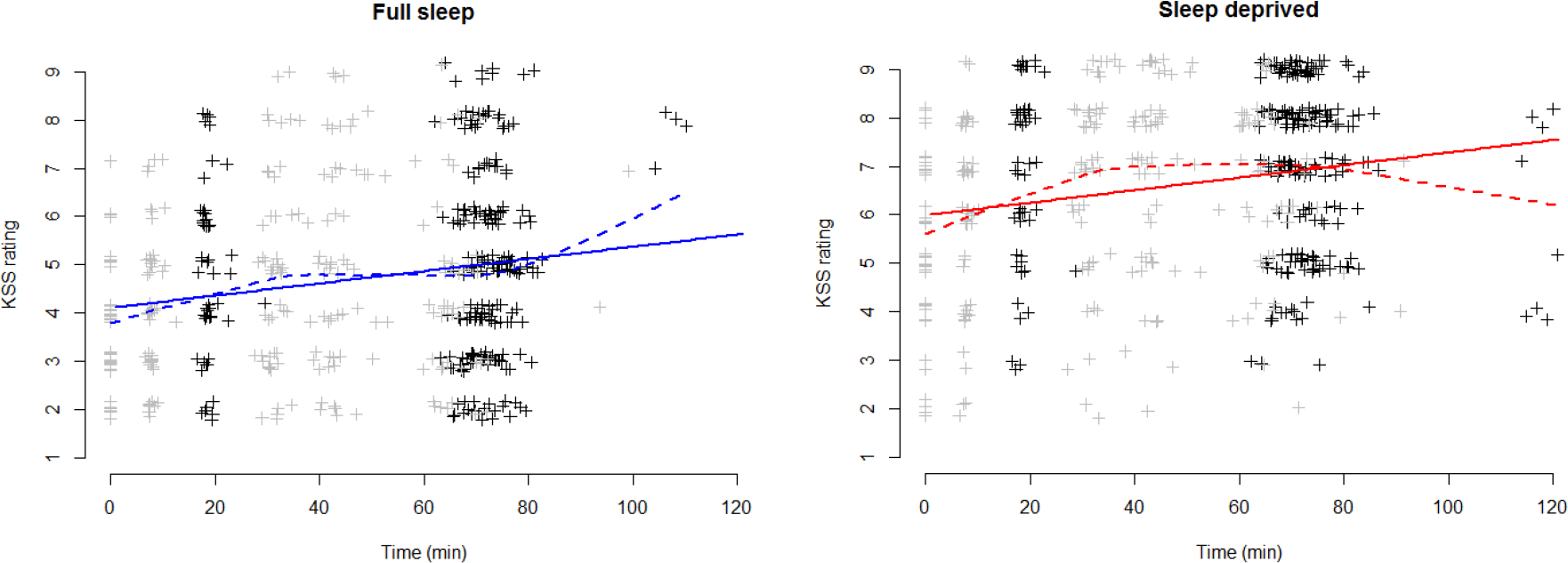
KSS ratings. Left: Full sleep condition. Right: Sleep deprivation condition. Data have been vertically jittered to aid visualisation. Points in black show KSS ratings made in connection to resting state experiments, and points in gray show other KSS ratings. Straight lines show linear regressions; dashed lines show loess regressions, for visualisation. In most cases, imaging was completed within 90 minutes and data points occurring later are mostly due to technical or other interruptions in the experiment, with participants exiting the MRI scanner and entering again.

### Head motion

We analysed head motion after scrubbing to verify that it was not a major confounder in comparisons between conditions and age groups. The results have been reported for the purpose of technical validation in Nilsonne et al.^56^. Briefly, sleep deprivation did not cause considerably more volumes to fall above the threshold for exclusion of framewise displacement > 0.5 mm in either run (estimates ≤ 1.2, p:s ≥ 0.2). However, more volumes exceeded the threshold in the second run compared to the first in both sleep conditions (estimates ≤ 3.8, *p*:s ≤ 0.0001). Older participants did not have more volumes exceeding the threshold in either condition nor session (estimates ≤ 1.0, p:s ≥ 0.35). These analyses suggest that head motion was not a major confounder between sleep deprivation and full sleep conditions nor between age groups.

### Independent component analyses

We performed 2 sets of ICA analyses; first one with 20 components^56^, using temporal filtering in the preprocessing and a gray matter mask, and secondly one with 30 components, without temporal filtering and without the gray matter mask. The two approaches yielded highly similar network results (supplemental Figure 2). The 30-component analysis yielded 10 components of interest, which were further examined in 2^nd^ level modelling for effects of sleep deprivation and differences between age groups. Network components of interest are shown in Figure 2. Network components not of interest are shown in supplemental Figure 2.

**Figure 2:**
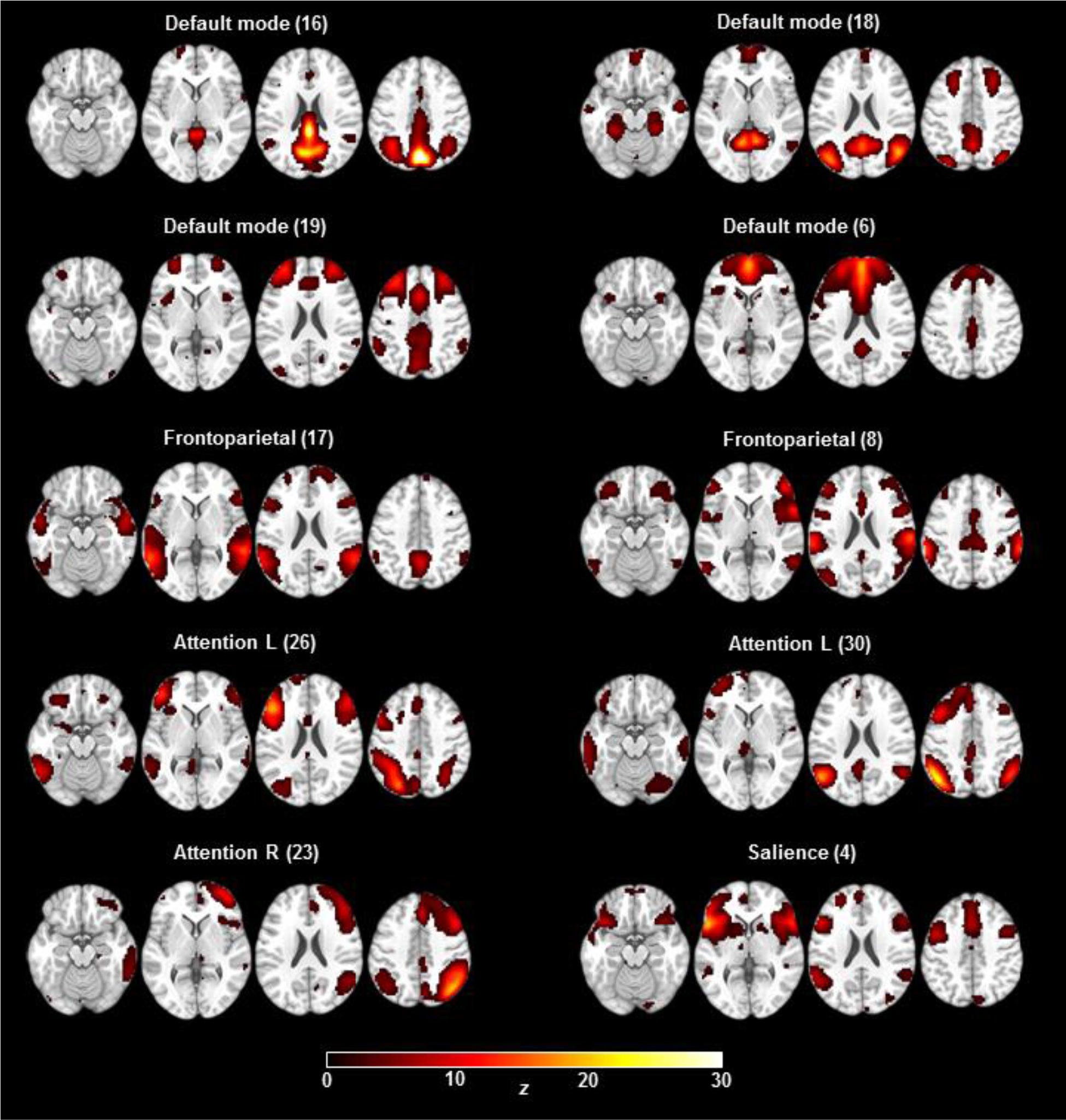
ICA networks of interest. Ten of 30 components (arbitrary numbers) were judged to represent the default mode network (DMN), frontoparietal, attention (lateralised left and right), and salience networks, and were examined further. Images are displayed in neurological convention. Scale is truncated at *z* = 30 and maps are thresholded at *z* = 1.645.

Sleep deprivation did not cause changes in connectivity within the networks of interest (*p_FWE_* < 0.05). Younger participants showed a pattern of greater connectivity than older participants within all networks (Figure 3). Older participants showed higher connectivity in small scattered foci (not shown; available at http://doi.org/10.5281/zenodo.581250). Sleep deprivation and age group showed no significant interactions.

**Figure 3:**
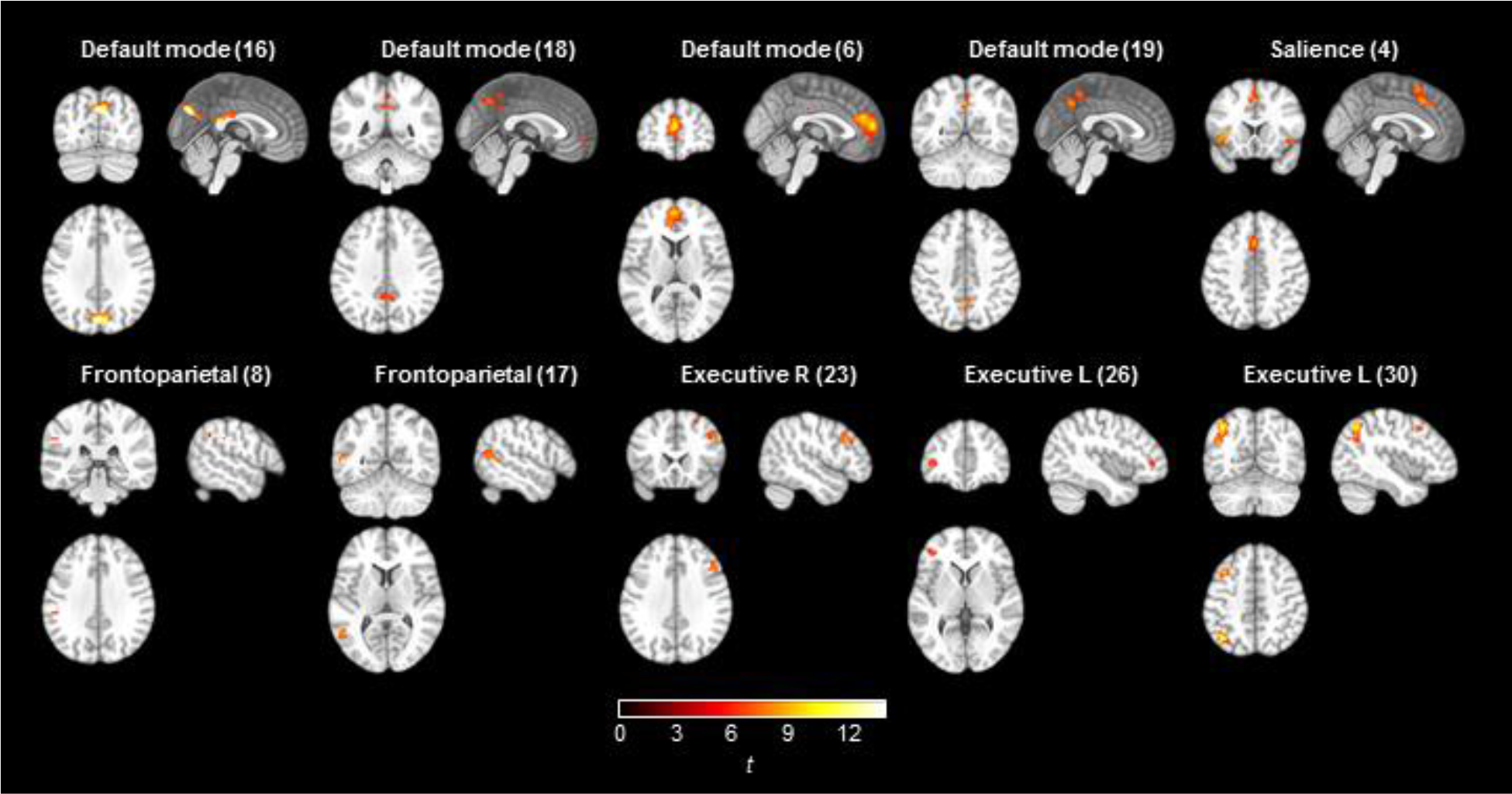
ICA results. Areas within network masks where younger participants showed higher functional connectivity compared to older participants (*p_FWE_* < 0.005, accounting for multiple comparisons; critical *t* = 5.45).

### Cross-correlation analyses of default mode and task-positive networks

Besides the data-driven ICA approach, we also tested a specific hypothesis-driven set of correlations, namely between the default mode network (DMN) and the task-positive network (TPN). We used seed regions defined in earlier work on sleep deprivation^10^ (Figure 4).

**Figure 4:**
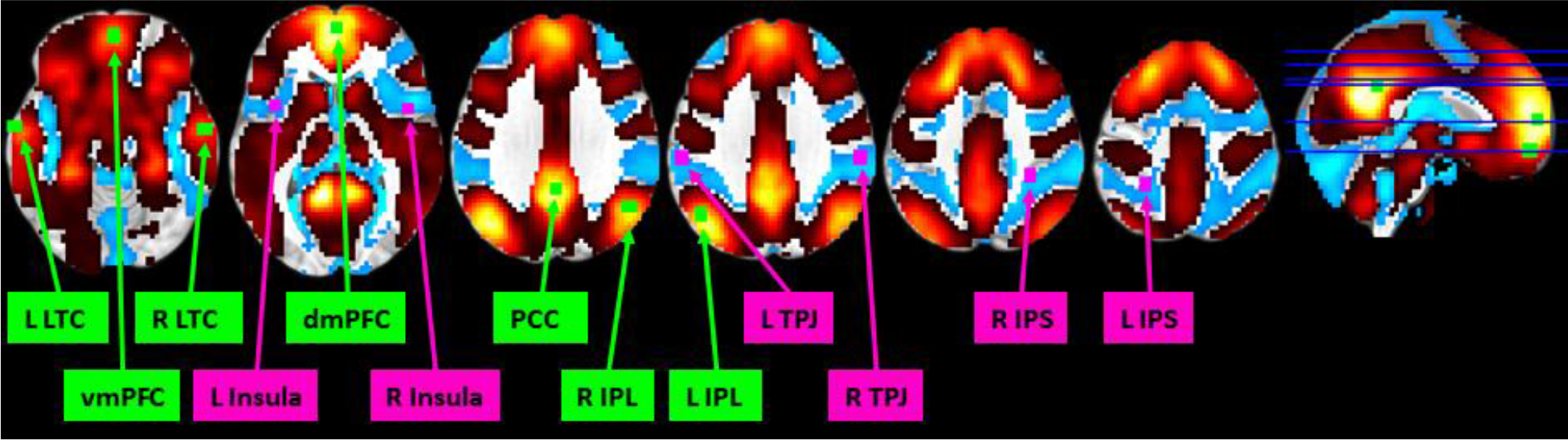
Seed regions for the default mode network (DMN, green) and task-positive network (TPN, purple), defined from de Havas et al.^10^. To verify that the seed regions are appropriate for the present dataset, they are superimposed on t-maps for positive connectivity (hot colors) and negative connectivity (cool colors) to regions from ICA components 18 and 19, representing posterior and anterior DMN (see methods and supplementary figures 7 and 8). IPL: inferior parietal lobule. IPS: inferior parietal sulcus. LTC: Lower temporal cortex. PCC: posterior cingulate cortex. dmPFC: dorsomedial prefrontal cortex. vmPFC: ventromedial prefrontal cortex. TPJ: temporo-parietal junction.

An overall pattern of correlation between DMN pairs and anticorrelation between DMN nodes and ACN nodes was confirmed (Figure 5). Sleep deprivation caused no changes surviving multiple comparisons correction at *p_FDR_* < 0.05, although the pattern of effects was in the expected direction, with lower connectivity within the default mode network and reduced anticorrelation between the default mode network and the task-positive network (Figure 5). Older participants showed a pattern of generally lower connectivity, with three default mode network node pairs surviving multiple comparisons correction at *p_FDR_* < 0.05 (Figure 5). For the interaction between sleep deprivation and older age, a pattern was observed where older participants showed less of the reduced connectivity suggested by the sleep deprivation main effects, although again no node pairs surviving multiple comparisons correction at *p_FDR_* < 0.05 (Figure 5).

**Figure 5:**
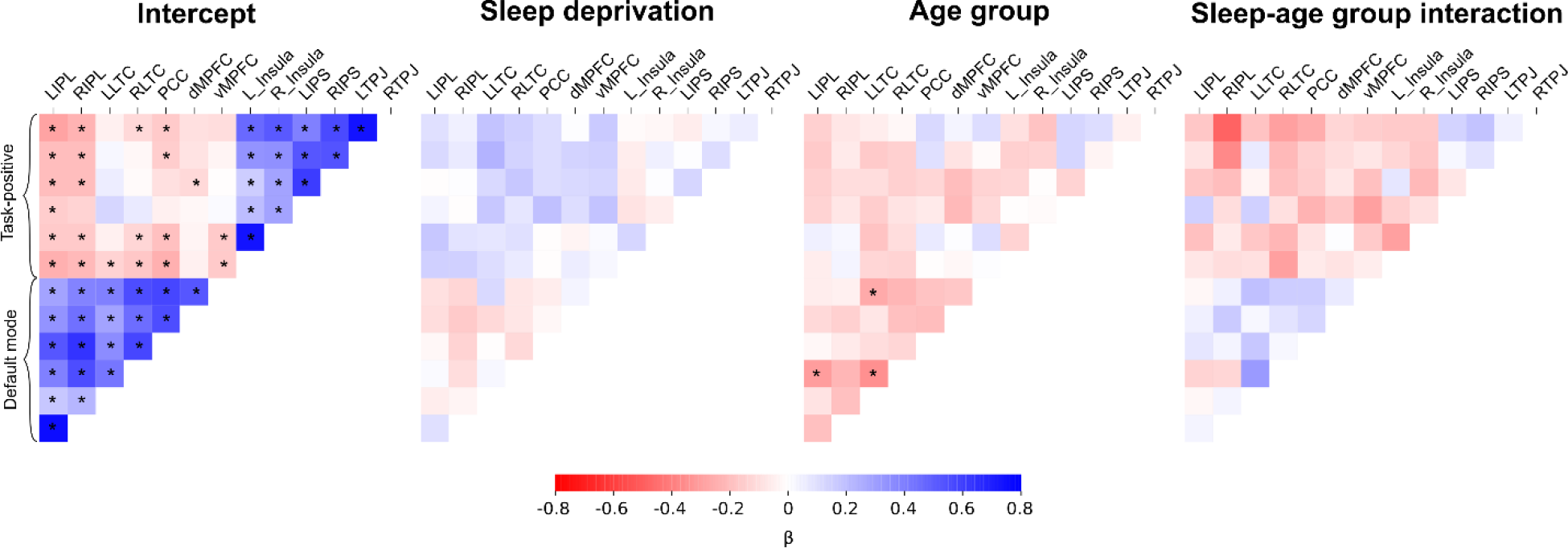
Connectivity results between regions of interest in the default mode network and the task-positive network. Associations between time courses from each ROI were determined for each run and entered into a mixed-effects model. Thus, the intercept represents the overall association between nodes; sleep deprivation represents the main effect of sleep deprivation vs full sleep; age group represents the main effect of older vs younger age group membership; and the sleep-age group interaction represents the interaction effect of sleep deprivation and older age. β is the standardized regression coefficient. False discovery rate correction was performed within each of the four sets of comparisons, and node pairs where *p_FDR_* < 0.05 are indicated with an asterisk (*).

Total sleep time was associated with increased connectivity between RIPL and LLTC in the interaction between older group and sleep deprivation (0.027, *p_FDR_* = 0.04). None of the other putative sleep-related covariates listed in the hypotheses (above) correlated significantly to ROI-based connectivity measures at *p* < 0.05 after false discovery rate correction for multiple testing within each contrast (main effect, interaction with sleep condition, interaction with age group, and 3-way interaction).

### Seed-based analyses of connectivity with thalamus and amygdala, and within the default mode network

Based on reports of effects of sleep deprivation on connectivity from the thalamus^12,22^ and amygdala^13,14^, we investigated whether earlier results could be replicated. Seed regions are shown in supplemental figures 3 and 4 for thalamus and amygdala, respectively. Functional connectivity with the thalamus was exhibited in the anterior and posterior cingulate cortices, the occipital cortex, and the cerebellum (supplemental Figure 5). Functional connectivity with bilateral amygdalae was shown in large parts of the brain, including contralateral amygdala, basal ganglia, and cortical areas in all four lobes (supplemental Figure 6). Sleep deprivation caused no differences in connectivity with thalamus or amygdala seeds exceeding a threshold of *p_FWE_* = 0.05, nor did any differences between the age groups exceed that threshold.

**Figure 6:**
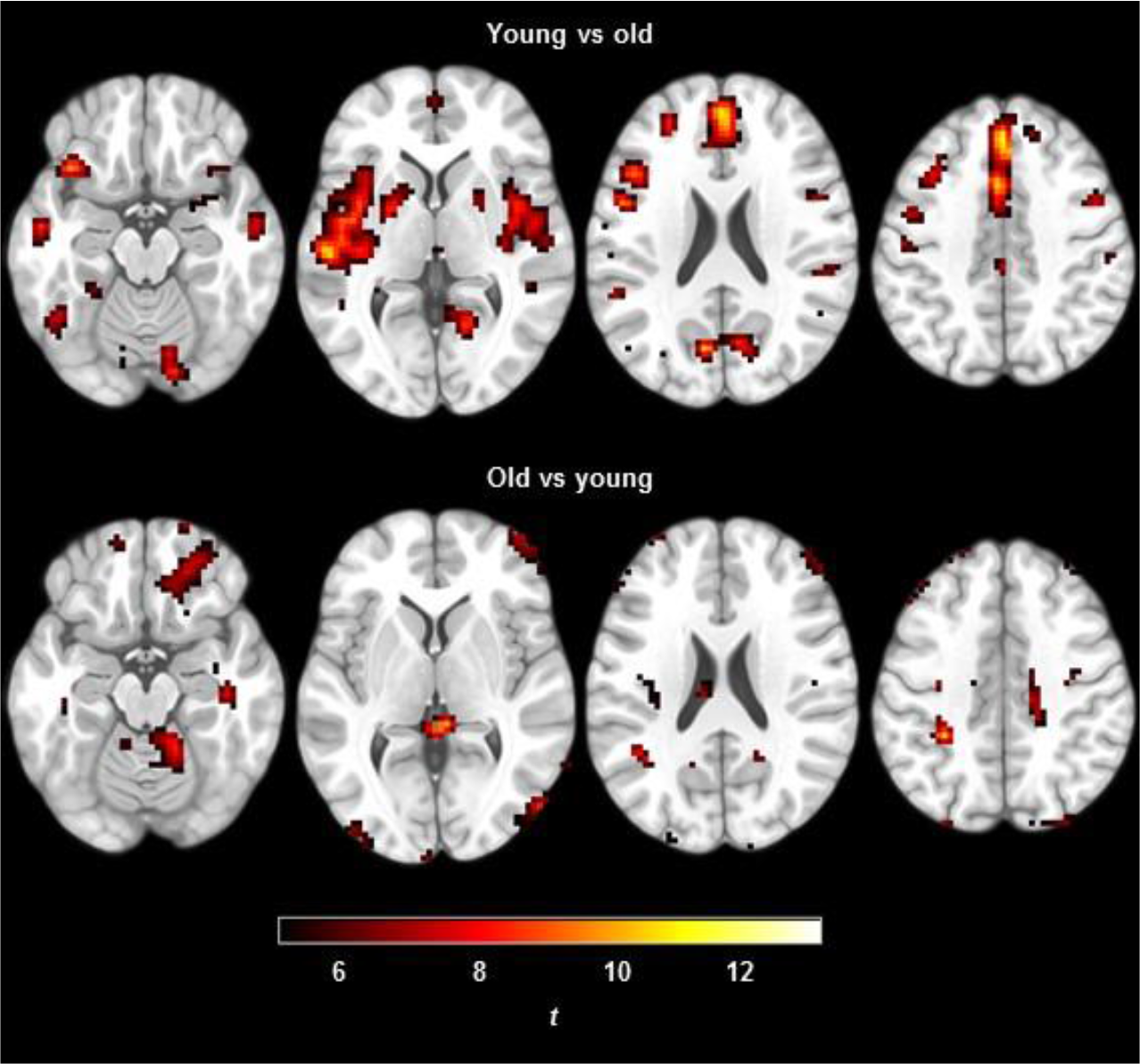
Differences in regional homogeneity between younger and older participants.

Following the suggestion of one reviewer, we investigated connectivity within the default mode network using seed regions defined from the ICA analysis results in the present dataset. The rationale was that this approach would be more sensitive to differences between conditions, albeit at increased risk of overfitting leading to reduced external validity. We therefore investigated connectivity from one seed region centered on the peak voxel in the posterior midline default mode network hub, as well as a seed region set defined by those areas where ICA-derived default mode network components showed *z* > 10. Resulting connectivity maps captured key areas of the default mode network (supplemental Figure 7). No effects of sleep deprivation, age group, nor their interactions were statistically significant. Negative connectivity maps from these seeds failed to show expected anticorrelated networks (supplemental Figure 7).

**Figure 7:**
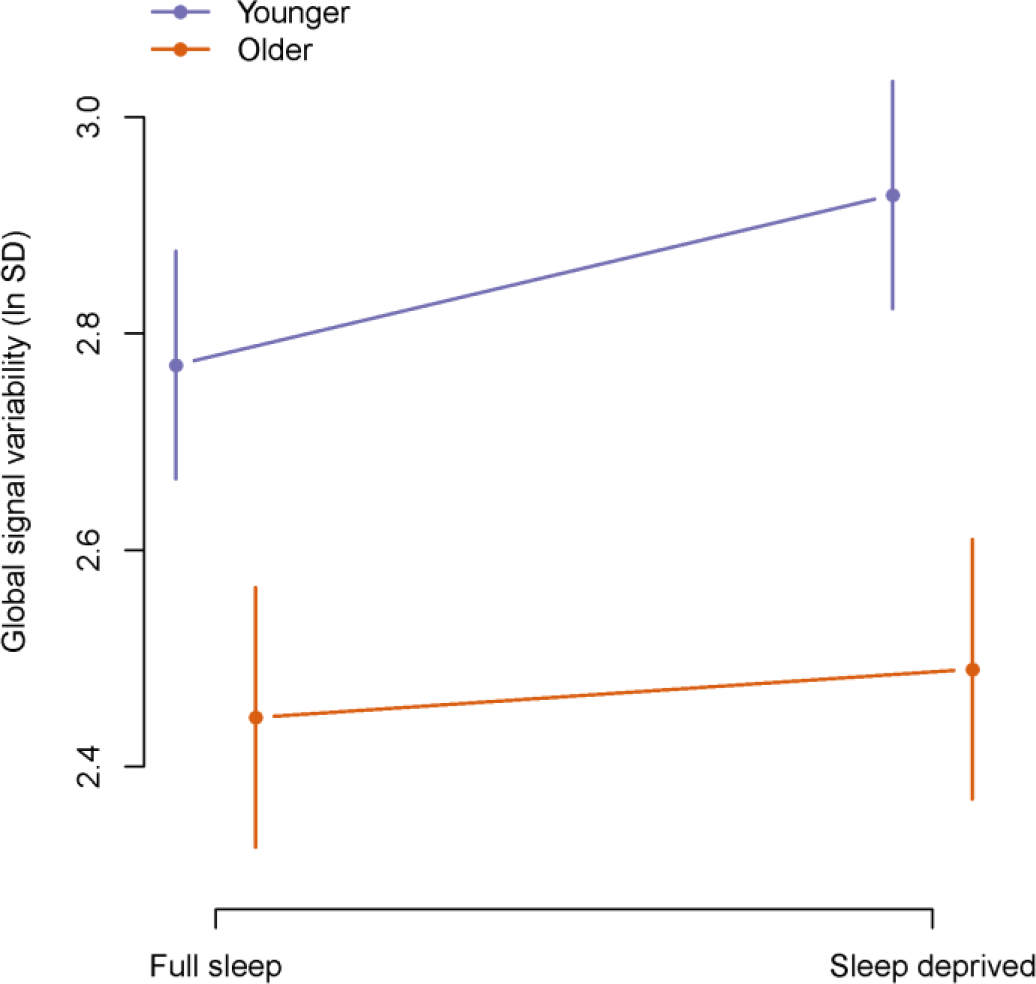
Global signal variability (standard deviation, transformed using the natural logarithm) after sleep deprivation in younger and older participants. Error bars show 95% confidence intervals.

### Regional homogeneity (ReHo)

Sleep deprivation did not affect measures of ReHo. Younger participants had higher ReHo in areas of the cerebral cortex and basal ganglia, particularly in the medial prefrontal cortex and in the superior temporal cortex/insula bilaterally (Figure 6). Older participants had higher ReHo mainly in areas prone to imaging and motion artifacts, including the orbitofrontal cortex and the outer edges of the brain anteriorly and posteriorly. Therefore, and even though framewise displacement was included as a 2^nd^-level regressor of no interest to account for motion artifacts, areas with apparently higher ReHo among the elderly may not reflect genuine differences in homogeneity of cerebral blood flow. There was no significant interaction between age group and sleep condition.

### Global signal

Global signal was not used as a covariate in our analyses reported above, since global signal regression has been identified as a potential source of spurious correlations^38,40^. However, we separately investigated whether sleep deprivation affected the global signal. Our main outcome of interest was global signal variability.

We found that sleep deprivation caused higher global signal variability (0.10 [0.03, 0.17], *p* = 0.003, Figure 7). Global signal variability was lower in older participants (−0.38 [-0.53, −0.23], *p* < 0.0001). The interaction between sleep deprivation and older age was −0.11 [−0.24, 0.02], *p* = 0.09. We found no notable associations between 15 putative predictors and global signal variability (supplemental Table 2), except that longer total sleep time (TST) in the sleep deprived condition was associated with less global signal variability (−0.01 [−0.01, 0.00], *p* = 0.01).

Effects of sleep deprivation and age group on the global signal itself, rather than its variability, are shown in supplemental Figure 9. Sleep deprivation and older age group interacted (106.0 [46.0, 166.0], *p* = 0.0006). The main effect of sleep deprivation was −15.1 [−45.1, 14.9], *p* = 0.32. The main effect of older age group was 44.2 [−87.3, 175.8], *p* = 0.50. We found no strong associations between 15 putative predictors and global signal (supplemental Table 3).

### Amplitude of low-frequency fluctuations

ALFF and fALFF were investigated exploratively as differences in ALFF following sleep deprivation were shown in a recent publication^24^. We found no significant differences between sleep conditions exceeding a threshold of *p_FWE_* > 0.05, and only scattered foci in comparisons between age groups (results not shown; available at http://neurovault.org/collections/LEWNNZLY/).

## Discussion

We found that partial sleep deprivation caused higher global signal variability. We did not find other major effects of partial sleep deprivation on measures of intrinsic connectivity, including ICA-derived networks; seed-based connectivity in the default mode and task-positive networks, and from the thalamus and amygdala to the rest of the brain; ALFF; and ReHo. Older participants generally showed less functional connectivity than younger participants. Major interactions between age group and sleep deprivation were not observed.

Our finding of increased global signal variability has been previously reported in one experiment with sleep deprivation, but without inferential statistics^22^. It is possible that global signal variability increased in sleepiness because of the increased propensity to drift in and out of sleep (wake-state instability), as the transition to sleep is associated with changes in network connectivity^74^. Our finding of increased global signal variability after sleep deprivation agrees well with earlier findings that global signal variability is decreased by caffeine^47^ and that global signal variability correlates to EEG measures of vigilance^48,49^. This finding is also consistent with findings by Fukunaga et al.^50^ that global signal variability increased in sleep, and by Kiviniemi et al.^75^ that midazolam sedation caused increased global signal variability. Further supporting a link between vigilance state and intrinsic connectivity fluctuatiuons, Wang et al.^76^ reported that dynamic functional connectivity analysis identified networks more active in low and high wakefulness, as determined by eye closures. Similarly, Chang et al.^77^ identified resting-state networks associated to eye closures in monkeys and found that fluctuations in these networks were related to cortical electrophysiological indices of arousal. Power et al.^78^ reported that effects of respiration, heart rate, and movement were associated to global signal variability even after nuisance regression. These findings suggest that wake-state instability causes variation in respiration and heart rate, which in turn affect the global signal. Under this interpretation, changes in physiological parameters of respiration and heart rate are mechanisms rather that confounders in the putative association between wakefulness and global signal variability. Further research using concomitant EEG and physiological recordings during resting state in humans may be able to shed light on this hypothesis.

We attempted replications of the analyses performed in Sämann et al.^8^ and in de Havas et al.^10^. In cross-correlation analyses of the default mode network and the task-positive network, the pattern of effect directions was consistent with results reported by de Havas et al.^10^, showing mainly reduced connectivity within the DMN and reduced anticorrelation between the DMN and the TPN. Our observed effects were however smaller in magnitude compared to those observed by de Havas et al. Against the backdrop of a number of earlier studies with consistent results, not all of which reached statistical significance, we view the results of our cross-correlation analysis as providing additional support for reduced connectivity within the DMN, and reduced anticorrelation between the DMN and the TPN, following sleep deprivation^29^.

An important difference between our study and several earlier studies is that we acquired resting state data with eyes open and with eye-tracking to ensure participants did not fall asleep^29^. If effects in earlier studies were due partly to sleep episodes during scanning, this difference in methodology may account for the weaker effects observed here compared to certain earlier reports.

In line with previous research^37^, older participants had lower connectivity within most ICA-derived networks of interest. Furthermore, we found that older participants had lower regional homogeneity (ReHo) in medial prefrontal cortex as well as superior temporal lobes and insula bilaterally. To our knowledge, only one study has previously investigated the effect of normal aging on ReHo in the resting brain, finding lower ReHo in motor areas^79^.

In the present study, we used a partial sleep deprivation paradigm, mainly because it has higher ecological validity compared with total sleep deprivation. Increased subjective sleepiness in the PSD condition confirms that the current protocol successfully induced sleepiness. The KSS measure of subjective sleepiness has been closely related to behavioral and physiological sleepiness^30^. KSS levels reached after PSD correspond to those seen during night work or night driving, although not as high as that seen before driving off the road in a driving simulator or being taken off the road for sleep related dangerous driving^30^. Thus, it is possible that the sleep manipulation might not have been strong enough to cause alterations in intrinsic connectivity measures, for which we found no effects. Thus, one possible conclusion is that partial sleep deprivation and the associated moderate sleepiness may not cause changes in intrinsic connectivity.

Limitations include the risk of confounding due to head motion, which is expected to cause an apparent decrease in long-range connectivity, e.g. within the default mode network^80^, and an increase in short-range connectivity, e.g. regional homogeneity. Although the number of excluded volumes due to head motion was not considerably greater in the sleep deprivation condition, and although we attempted to correct for motion by regressing out realignment parameters, we cannot exclude the possibility of residual effects. Another possible improvement would have been to acquire EEG during resting state scans. This could have allowed us to identify microsleeps. We do not believe that the wake up-regime used in this study (which was almost never activated) prevented microsleeps with eyes open from occurring. Strengths of this study include a relatively large sample that includes both younger and older adults, recording of PSG and KSS, and monitoring of participants by eye-tracking.

In conclusion, we report that global signal variability was increased by sleep deprivation. We speculate that this effects is due to wake-state instability, affecting neural activity as well as respiration, heart rate, and head movements. Major effects of sleep deprivation on resting state networks were not observed.

## Acknowledgements

We are grateful to Paolo d’Onofrio, Diana S. Cortes, Dani Cosme, and Roberta Nagai for expert assistance with polysomnography recordings, to Birgitta Mannerstedt Fogelfors for assistance with data acquisition, and to Rouslan Sitnikov and Jonathan Berrebi for expert assistance with MRI sequences and auxiliary equipment and stimulus presentation software.

This work was supported by Riksbankens Jubileumsfond, Fredrik and Ingrid Thuring’s Foundation, and the Karolinska Institutet Strategic Neuroscience Program.

## Authors’ contributions

Designed the study: GN, ST, JS, HF, GK, ML, PF, TÅ. Acquired data: GN, ST. Analysed data: GN, RA, PF. Interpreted results: GN, ST, JS, RA, HF, GK, ML, PF, TÅ. Drafted manuscript: GN. All authors read and approved the final version of the manuscript.

